# Analysis of miniaturization effects and channel selection strategies for EEG sensor networks with application to auditory attention detection

**DOI:** 10.1101/593194

**Authors:** Abhijith Mundanad Narayanan, Alexander Bertrand

## Abstract

**Objective:** Concealable, miniaturized electroencephalo-graphy (‘mini-EEG’) recording devices are crucial enablers towards long-term ambulatory EEG monitoring. However, the resulting miniaturization limits the inter-electrode distance and the scalp area that can be covered by a single device. The concept of wireless EEG sensor networks (WESNs) attempts to overcome this limitation by placing a multitude of these mini-EEG devices at various scalp locations. We investigate whether optimizing the WESN topology can compensate for miniaturization effects in an auditory attention detection (AAD) paradigm.

**Methods:** Starting from standard full-cap high-density EEG data, we emulate several candidate mini-EEG sensor nodes which locally collect EEG data with embedded electrodes separated by short distances. We propose a greedy group-utility based channel selection strategy to select a subset of these candidate nodes, to form a WESN. We compare the AAD performance of this WESN with the performance obtained using long-distance EEG recordings.

**Results:** The AAD performance using short-distance EEG measurements is comparable to using an equal number of long-distance EEG measurements if in both cases the optimal electrode positions are selected. A significant increase in performance was found when using nodes with three electrodes over nodes with two electrodes.

**Conclusion:** When the nodes are optimally placed, WESNs do not significantly suffer from EEG miniaturization effects in the case of AAD.

**Significance:** WESN-like platforms allow to achieve similar AAD performance as with long-distance EEG recordings, while adhering to the stringent miniaturization constraints for ambulatory EEG. Their applicability in an AAD task is important for the design of neuro-steered auditory prostheses.

## I. Introduction

Electroencephalography (EEG) is a non-invasive neurorecording technique, which has the potential to be used for 24/7 neuromonitoring in daily life, e.g., in the context of neural prostheses, brain-computer interfaces, or for improved diagnosis of brain disorders [2]–[17]. Although existing mobile wireless EEG headsets are a useful tool for short-term experiments, they are still too heavy, bulky and obtrusive, for long-term EEG-monitoring in daily life. However, we are now witnessing a wave of new miniature EEG sensor devices containing small electrodes embedded in them, which we refer to as mini-EEG devices. These mini-EEG devices are concealable and light-weight, and come in various forms. Among these forms, some can be concealed behind the ear [5], [11], [12], placed in the ear [6], [7], [15], [18], attached to the skin as a stick-on tattoo [4] or inserted under the skin [3]. The research towards novel concealable mini-EEG devices is still highly active, with regular emergence of new innovative form factors.

However, due to their miniaturization, these mini-EEG devices have the drawback that only a few EEG channels can be recorded within a small area. Therefore, to capture more spatial information, one could use a multitude of such devices and wirelessly connect them in a sensor network-like architecture, referred to as a wireless EEG sensor network (WESN) [13], [19]. The EEG measured in such a WESN will consist of local short-distance measurements made by multiple mini-EEG devices or ‘nodes’, which consist of at least two electrodes. This is unlike EEG recordings made by traditional headsets where electrodes are typically referenced to a common reference electrode or an average reference signal. In this paper, we carry out a comprehensive study on the effect of short-distance measurements recorded by the nodes of a WESN and we propose a method to find the optimal scalp locations to place those nodes. The nodes of this WESN are emulated by re-referencing standard cap-EEG electrodes to nearby electrodes.

In this work, we consider the application of auditory attention detection (AAD) to explore the impact of these short-distance measurements. Several studies have successfully demonstrated that it is possible to estimate the attended speech envelope from EEG [8], [16], [20]–[24], thereby detecting which speaker a subject is attending to in a multi-speaker listening environment (the so-called ‘cocktail party’ problem). It is believed that in the future these AAD systems can be used for the cognitive control of auditory prostheses, such as hearing aids and cochlear implants [23], [24]. Therefore, AAD is an application that could benefit hugely from chronic neuromonitoring using a WESN-like platform.

We compare the AAD performance between the short-distance EEG measurements acquired by single-channel (two-electrode) mini-EEG nodes and two different long-distance EEG measurement configurations. In the first long-distance configuration, EEG measurements were obtained from electrodes with respect to a common reference electrode (Cz). In the second configuration, EEG measurements were obtained with the freedom to choose any electrode-reference pair. The comparison between short and long-distance configurations was carried out after selecting the best subset of channels in each case, which equates to choosing the most relevant positions on the scalp. To this end, we use a greedy utility-based sensor subset selection method [25], [26] to find these optimal locations on the scalp. We verify the suitability of this method in the channel selection problem for AAD by comparing it to two other methods in literature, viz. a decoder magnitude-based method [20] and the group-LASSO (least absolute shrinkage and selection operator) based channel selection method which was used in [1]. We determine the optimal locations on the scalp in both a subject-dependent and subject-independent scenario. In the current work, we demonstrate that, when placed on these optimal locations, the short-distance EEG measurements acquired by a multitude of single-channel mini-EEG nodes do not significantly affect the AAD performance compared to both long-distance EEG measurement configurations. In addition, we also investigate the effect of adding a third electrode to each node used in the above experiments, to study the impact of having an additional channel. We show that this additional channel in each node, increases the AAD performance significantly, but only if the decoding weights can adapt to the individual subject. We also show that the optimal locations for both single-channel and two-channel nodes correspond to the temporal lobe regions associated with the auditory cortex, as reported in other experiments on EEG channel selection for AAD [20].

In the sequel, we will consistently use the following terminology:

- *Channel*: A channel is an EEG signal that originates from a single electrode pair over which the scalp potential is measured. Every channel has a one-to-one correspondence with a pair of electrodes. We will generally make abstraction of the ambiguity in polarity, as we work with data-driven methods which are not affected by the latter.
- *Node*: A node represents a group of (at least two) closely spaced EEG electrodes such as included in a wireless mini-EEG sensor device, emulated here as a group of nearby cap-EEG electrodes. One of these electrodes is treated as a reference electrode, which forms pairs with all the other electrodes in the node. As such, a node with *N* electrodes will contain *N* − 1 EEG channels. In this paper, we discuss single-channel and two-channel nodes which consist of two and three electrodes respectively.

The paper is organized as follows. In Section II, first the EEG data collection and AAD algorithm based on a least-squares (LS) estimation is explained followed by the details on the WESN emulation. This section also describes the channel selection strategies that are used in our comparison. In Section III, we show results on the AAD performance in a WESN setting and benchmark it against other recording settings with long-distance reference electrodes. Here, we also show the impact and possible benefits of adding an extra channel to each EEG node. Discussions on the results are given in Section IV and conclusions are drawn in Section V.

## II. Methods

### A. Experiment Data Collection

This paper reports experiments carried out using the data set described in [22]. The data set contains 16 subjects who listened to two simultaneous speakers coming from two distinct spatial locations, and were asked to attend to only one of them while ignoring the other. Half of the speech stimuli were presented to the subjects dichotically, while the other half was presented using head-related transfer functions to simulate a realistic acoustic scenario. The side of attention (left or right) was evenly split over the different trials to avoid decoder bias [27]. During the entire experiment, 64-channel EEG was recorded using a BioSemi ActiveTwo system resulting in 72 minutes of EEG data per subject. The electrodes were placed on the head according to international 10-20 standards and data was recorded with a common reference montage, with the Cz electrode used as the reference.

### B. Auditory Attention Detection

In this subsection the basic AAD procedure that is used in this paper is reviewed. In [8] it was shown that AAD can be achieved by a least-squares (LS) based reconstruction of the attended speaker’s speech envelope using multi-channel EEG recordings. Assuming the EEG data is split into *Q* trials of equal length, the goal is to detect for each trial to whom of the two speaker the subject was attending.

First, a linear spatio-temporal decoder ŵ that estimates the attended speech envelope from the *C*-channel EEG data is obtained by solving the following LS optimization problem:

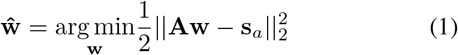

where **s**_*a*_ is a vector containing the attended speech envelope samples (which is assumed to be known during a training phase) and **A** is a matrix containing *M* copies of the *C* EEG channels in its columns (i.e., *M* ⋅ *C* columns in total), in which a delay of *j* − 1 samples is added to the *j*-th copy of each channel. We selected *M* = 5 in all of our experiments. The solution of Eq. (1) is given by

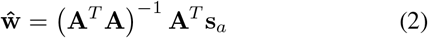

which can be written on a per-trial basis as

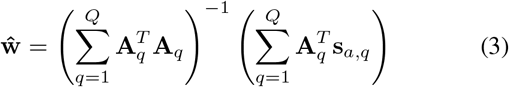

where **A**_*q*_ and **s**_*a,q*_ are the submatrices of **A** and **s**_*a*_ corresponding to trial *q*.

The experiments reported in this paper are based on trials of length 60s. To exclude over-fitting effects in the validation procedure, we use a subject-specific leave-one-out cross-validation where each trial is used once as a test trial, while all the other trials are then used to train a decoder for the specific trial under test. In [8] the cross-validation is carried out by computing Eq. (1) over the data of individual trials, after which the resulting per-trial decoders are averaged across trials, except for the test trial *k*. Here, we use a modification to the above method as proposed in [22], which constructs the decoder ŵ_*k*_ to decode trial *k* by including the data from all the trials except trial *k* to construct the matrix **A** and the vector **s**_*a*_:

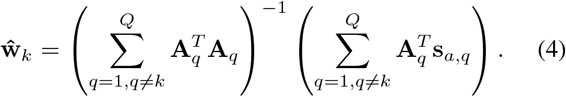

This procedure reduces or eliminates the need for regularization if sufficient data is available in the training set, or equivalently, if the number of rows in **A** is large enough [22].

Once the decoder ŵ_*k*_ is obtained, an estimate of the attended speech envelope in trial *k* is constructed using:

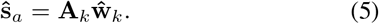

The attended speech envelope reconstruction is followed by the Pearson correlation coefficient computation between the reconstructed envelope ŝ_*a*_ and the two speakers’ speech envelopes. Therefore two correlation coefficients, *r*_*a*_ and *r*_*u*_ are calculated for each trial, where *r*_*a*_ is the correlation between the reconstructed speech envelope ŝ_*a*_and the attended speech envelope and *r*_*u*_ is the correlation between ŝ_*a*_and the unattended speech envelope. A trial is correctly decoded if *r*_*a*_ > *r*_*u*_ and the percentage of trials successfully decoded is used as the AAD performance parameter.

### C. WESN Emulation

The aim of this paper is to understand the effect of using short inter-electrode distances between the electrodes in order to emulate a WESN with the different nodes representing different mini-EEG devices. This emulation is achieved by re-referencing 64 cap-EEG channels towards nearby reference electrodes generating a set of candidate node locations and orientations.

First, we emulated a WESN in which each node consists of two electrodes separated by a short distance. Since each node then corresponds to a single electrode-pair, we refer to them as single-channel nodes. These nodes are selected from a set of candidate nodes created by pairing each electrode of the 64-electrode cap with each of its nearby electrodes that are at a distance of at most 5 cm. The distance was selected to ensure that a large number of candidate node locations and orientations are generated but at the same time the electrodes of each node have a reasonably short distance between them to emulate a miniaturized EEG-sensor node. Using this criteria, a set of *P* = 209 candidate single-channel node locations and orientations were generated from the original 64 electrodes with an average inter-electrode distance of 3.7 cm. We will refer this set as *S*_1*ch*_. It is noted that *P* > 64, hence these *P* EEG channels form a redundant (linearly independent) set.

Second, we emulated a WESN with nodes containing three electrodes with one of them acting as a reference electrode, resulting in two EEG channels per node. In [17], it was shown for behind-the-ear electrodes that a smaller angle between two electrode pairs leads to a higher correlation between the recorded signals at both pairs. Therefore, we ensured that the two corresponding electrode pairs in the three-electrode node have a near-orthogonal orientation, based on the following procedure. For each electrode *k* = 1, …, 64, we again select all the electrodes that are within 5 cm distance of electrode *k*, forming a set of candidate electrode pairs denoted by *P*_*k*_. For each electrode pair *p* ∈ *P*_*k*_, we select all the pairs *q* ∈ *P*_*k*_\{*p*} for which the angle between pair *p* and pair *q* is between 60 and 120 degrees in the 3-D coordinate space. All these combined pairs *{p, q}* form a new set *C*_*k*_ containing all candidate 2-channel nodes which have electrode *k* as the reference, where the duplicates are removed. The total pool ∪_*k*_*C*_*k*_ contains *P* = 203 candidate 2-channel nodes. This set of 2-channel nodes will be referred as *S*_2*ch*_. Note that *P* refers to the number of nodes, and since each node can have more than one channel, the total number of channels is larger or equal to *P*. Let the total number of EEG channels across all candidate nodes be *P′* (channels that belong to multiple nodes are also counted more than once). For *S*_1*ch*_, *P′* = *P* and for *S*_2*ch*_, *P′* = 2*P*.

### D. Long-distance EEG measurement benchmark

To study the impact of short-distance EEG measurements, we created two long-distance benchmark EEG measurement sets. First is the original EEG measurement where each electrode is referenced to the Cz electrode. We refer to this set as the ‘Orig (Cz-ref)’ set of EEG measurements. However, although the Orig (Cz-ref) case allows recordings over larger distances than the emulated WESN, it may also be penalized in the sense that each of the N selected channels should use the same reference electrode, thereby reducing the choice of the orientation of the electrode pair. Therefore, we created a second benchmark set consisting of all possible electrode-pairs from the original EEG data without any constraints on the distance. This creates a total of *P* = 2016 candidate pairs. We refer to this set as the ‘Any-Pair’ set of EEG measurements.

### E. Decoder magnitude-based (DMB) node selection

To construct a WESN, the main objective is to select the *N* best nodes from *P* node candidates (we will consider the case of single-channel nodes (*S*_1*ch*_) in Section III-B1 and the case of two-channel nodes (*S*_2*ch*_) in Section III-B2). To this end, we replace **A** in Eq. (1), with **A**_*P*_ which now contains the *P ^l^* EEG channels across all *P* candidate nodes. Note that the same channel can appear multiple times in **A**_*P*_ if that channel is included in more than one node. Therefore the optimization problem in Eq. (1) changes to:

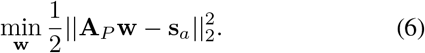

where **w** is now a (*P′* ⋅ *M*)-dimensional vector variable. It is noted that the matrix **A**_*P*_ is rank deficient as *P′* > 64, i.e., there are many linear dependencies between the channels of different nodes in the candidate sets *S*_1*ch*_ and *S*_2*ch*_, which means that there are infinitely many solutions for Eq. (6). This problem is typically solved by selecting the solution with minimal *l*_2_-norm, which is known to be beneficial to reduce overfitting effects [26]. The minimum-norm solution of Eq. (6) is given by

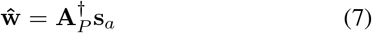

where 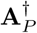 denotes the Moore-Penrose pseudo-inverse of **A**_*P*_, which can be computed as

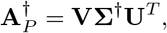

where

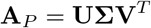

is the singular value decomposition of **A**_*P*_, and where **Σ**^*†*^ is formed by taking the reciprocal of each non-zero element on the diagonal of **Σ**, and then transposing the matrix [28].

In [20] and [29], an iterative channel selection heuristic was used for EEG-based AAD, in which the channel with the lowest corresponding entry in the decoder **ŵ** was removed in each iteration. We will refer to this channel selection method as the decoder magnitude-based (DMB) method, which is here applied to the minimum-norm solution given in Eq. (7).

### F. Greedy utility-based node selection

In general, the weights in an LS or a linear minimum mean squared error (MMSE) decoder do not necessarily reflect the importance of the corresponding channels to minimize the squared error cost^1^[26]. As such, the metric used in the DMB node selection process does not truly capture the contribution of each channel to the least squares envelope reconstruction. Therefore, we propose the use of the so-called ‘utility’ metric instead to perform a greedy channel selection, which we benchmark against the DMB method and another commonly used variable selection algorithm known as LASSO (see Section II-G).

For the sake of an easy exposition, we will first explain how to use the utility metric to greedily select *N columns* of **A** or **A**_*P*_. We will later explain how this can be extended to the selection of *nodes*, i.e., at the granularity of pre-defined groups of columns rather than individual columns. To select the subset with the *N* ‘best’ columns of **A**, we use an iterative greedy method based on the utility metric to eliminate columns one by one. In the context of an LS problem like Eq. (1), the utility of a column is defined as the increase in the squared error when the column would be removed, and the decoder **w** would be re-optimized for the remaining set of columns [25],[26]. Remarkably, the utility of each column can be monitored in an efficient way without the explicit recomputation of the optimal decoder for each column removal, which would imply a strong computational burden. By defining the inverted auto-correlation matrix

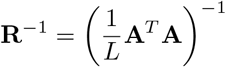

where *L* is the number of rows in **A**, the utility of the *k*-th column can be computed as [25],[26]:

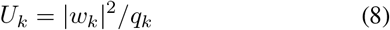

where *q*_*k*_ is the *k*-th diagonal element of **R**^−1^, and *w*_*k*_ is the *k*-th decoder weight of **ŵ** as defined in Eq. (2). Note that **R**^−1^ is immediately available from the calculation of Eq. (2), hence the utility of each column can be computed as a by-product of the calculation of Eq. (2).

However, the matrix **A**_*P*_ in Eq. (6) is rank deficient and therefore contains redundant columns that are linear combinations of the other columns. Therefore, removal of a redundant column from **A**_*P*_ will not lead to an increase in LS cost of Eq. (6). As all columns in the initial set are redundant, all columns would have zero utility by definition. Furthermore, **R**^−1^ will not exist in this case as the matrix **A**_*P*_ is rank deficient. To overcome this problem, we use the definition of utility generalized to a minimum *l*_2_-norm selection [26] which eliminates the redundant column yielding the smallest increase in the *l*_2_-norm of the decoder when that column were to be removed and the decoder would be re-optimized. As mentioned in Section II-E, minimizing the *l*_2_-norm of the decoder reduces the risk for overfitting. This generalization is achieved by first adding an *l*_2_-norm penalty to the cost function that is minimized in Eq. (6):

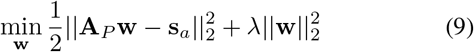

where 0 < *λ* ≪ *ϵ* with *ϵ* equal to the smallest non-zero squared singular value of **A**_*P*_. The minimizer of Eq. (9) is:

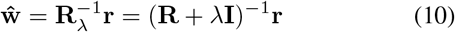

where 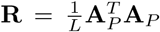 and 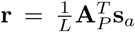, are referred to as the autocorrelation matrix and cross-correlation vector respectively. The utility *U*_*k*_ of the *k*-th column in **A**_*P*_ based on Eq. (9) is [26]:

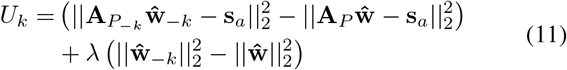

where **A**_*P−k*_ denotes the matrix **A**_*P*_ with the *k*-th column removed and **ŵ**_*−k*_ is the LS solution corresponding to **A**_*P−k*_. From Eq. (11), we see that, if the increase in LS cost is non-zero (i.e., column *k* is linearly independent from the other columns), then the first term dominates the second term, yielding the original definition of utility. However, if column *k* is linearly dependent, the first term vanishes and the second term will dominate. In this case, the utility quantifies the increase in *l*_2_-norm after removing the *k*-th column. Therefore, by iteratively removing the column with the lowest utility *U*_*k*_, we greedily reduce the number of columns while keeping both the squared error and the *l*_2_ norm small.

The greedy method of removing columns iteratively by computing their utility using Eq. (11) requires considerable computation time since we need to recompute the optimal decoder using Eq. (10) a number of times equal to the remaining number of columns in each iteration. This has an asymptotic computational complexity of *O*(*P′*^4^) [26], which is practically impossible for such large values of *P′* as those targeted in our experiments. However, similar to Eq. (8), it has been shown in [26] that *U*_*k*_ as defined in Eq. (11) again can be efficiently computed using Eq. (8). where *q*_*k*_ is now the *k*-th diagonal element of 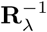 and *w*_*k*_ is the *k*-th element of **ŵ** as defined in Eq. (10). The use of Eq. (8) removes the need of multiple re-computations of optimal decoders in every iteration resulting in a reduction of computational complexity by three orders of magnitude.

As mentioned in Section II-E, **A**_*P*_ was constructed out of the EEG data from *P* nodes and their *M* − 1 delayed versions. Hence, the utility of a node would not just be the utility of a single column in **A**_*P*_ but of a group of columns, viz. a node’s channel(s) and its (their) delayed versions. To this end, we use the extension towards group-utility described in [26] and [30] which finds the utility of a group of columns. Consider, without loss of generality (w.l.o.g.), that **A**_*P*_ is formed using the set *S*_1*ch*_. Assume also w.l.o.g. that the node for which we calculate the utility, say node *p*, corresponds to the last *M* columns of **A**_*P*_, the node’s channel as well as its *M* − 1 delayed versions. Define the block-partitioning of 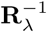:

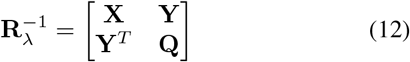

where, **Q** is an *M* × *M* matrix. Then, the group-utility of the columns of **A**_*P*_ corresponding to node *p* can be efficiently computed as [26]:

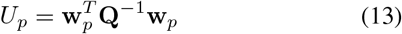

where **w**_*p*_ contains the last *M* entries of **ŵ**. Note that Eq. (13) reduces to the single-column utility of Eq. (8) if *M* = 1. The computation of group-utility using Eq. (13) is much faster than removing *M* columns of **A**_*P*_ and recomputing the optimal decoder. If the utility of all channels has to be computed multiple times, e.g., in a greedy selection procedure as described below, this allows to reduce the computation time from a few hours to a few seconds on a laptop with Intel Core i7-6820HQ clocked at 2.70GHz running Matlab R2015b.

To select *N* (out of *P*) nodes, we greedily remove the nodes with the lowest utility one by one as follows. First, the group-utility of each node’s channels (and their *M* − 1 delayed copies) are computed using Eq. (13) followed by the removal of the node with the least utility. After this removal, Eq. (10) is recomputed for the remaining set of nodes, after which the utility is again re-computed. This step is repeated until *N* nodes are left.

Finally, it is noted that the utility-based and the DMB greedy methods become equivalent in the case where the different channels are uncorrelated and have equal variance, in which case 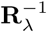 reduces to a scaled identity matrix. However, this is usually not the case in (high-density) EEG data.

### G. Group-LASSO based node selection

Another candidate algorithm which we have included in our channel selection benchmark is the commonly used LASSO-based variable selection method. LASSO adds an *l*_1_ norm penalty term to an LS regression problem like Eq. (6) to obtain a sparse solution for **ŵ**, i.e. a vector with few non-zero entries [31], where the non-zero entries in **ŵ** then correspond to the selected columns. However, our objective is to select groups of variables in **ŵ** that correspond to a particular node consisting of a set of channels and their delayed versions. Yuan and Lin [32] have proposed the group-LASSO (gLASSO) criterion to solve the aforementioned problem. gLASSO is a modification of LASSO for linear regression which introduces a sparse selection of pre-defined groups of variables without imposing sparsity *within* the individual groups. Applying the gLASSO criterion, Eq. (6) is modified as:

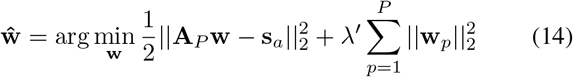

where 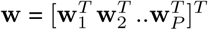, with **B**_*p*_ the sub-vector of length *M* of the decoder which contains the weights corresponding to the channels of node *p* and their *M* − 1 delayed versions, and where *λ′* is a tuning parameter which controls the sparsity of **ŵ**. Note that Eq. (14) has an *l*_1_-norm penalization across groups (represented by the summation sign), whereas each group is represented by the *l*_2_-norm over its coefficients. The optimal set of *N* channels can be found by increasing the parameter *λ′* until the gLASSO procedure selects exactly *N* channels.

### H. Subject-dependent vs subject-independent (universal) node selection

In our experiments, we performed a subject-dependent as well as a subject-independent channel selection or node selection. In the subject-dependent selection, **A**_*P*_ contains the *P′* EEG channels and their *M* − 1 delayed copies of subject *i* where *i* = 1, 2, *K* and *K* is the total number of subjects (*K* = 16 in this work). This results in a different decoder per subject, and hence a different selection of channels per subject. In the subject-independent selection, we use the data from all subjects, resulting in the stacked matrix:

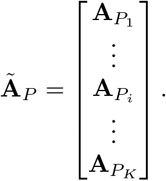

Here, **A**_*P*_*i*__ contains the *P′*–channel EEG data and their *M* − 1 delayed copies of subject *i*. By also replacing **s**_*a*_ in Eq. (6) with

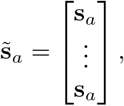

we can solve Eq. (6) to find a universal decoder that fits the data of all subjects.

Applying channel selection on this problem will then also result in a subject-independent selection of the best node locations. In the sequel, we will use the terms ‘universal decoder’ and ‘universal node selection’ to refer to the subject-independent decoding and subject-independent node selection, respectively.

Note that, after making a universal node selection, we have the option to either train a subject-dependent decoder or a universal decoder on the selected nodes. Both will be analyzed in the sequel, with leave-one-trial-out and leave-one-trial-and-subject-out cross-validation, respectively. In both cases, the prior node selection excludes the data from the subject under test. However, due to excessively large computation times, channel selection was not carried out in a leave-one-trial-out basis (although the trial is excluded when testing AAD performance).

## III. Results

### A. Benchmark of channel selection methods

Before investigating the miniaturization effects, we verified the suitability of the proposed utility-based greedy channel selection method on the problem of channel selection for the original EEG-cap data by comparing the proposed method with the DMB method used in [20] and the gLASSO method used in [1].

The three methods were used to select the *N* = 1, 2, 3*…*62 best (subject-dependent) channels on the original EEG data^2^. The AAD performance was computed for the channels selected for each value of *N*. The results are plotted in Fig. 1. When *N* > 25, the median AAD decoding accuracy remains more or less equal to the accuracy obtained with all channels, for all three channel selection methods. However, for *N* < 20 the median utility-based channel selection performance remains closer than the other methods to the baseline performance. Fig. 1 (b), which zooms in on AAD performances for *N* < 25, shows that the drop in performance occurs only at *N* < 10 for the utility-based method, while, for the DMB and gLASSO based method, this drop occurs earlier at *N* = 15. In addition, for small values of *N* the utility-based channel selection method is significantly better, than both the other methods. This was confirmed by performing a paired t-test and Wilcoxon signrank test between the decoding accuracies of the utility-based and DMB method on the one hand, and the utility-based and gLASSO method on the other hand.

**Fig. 1:**
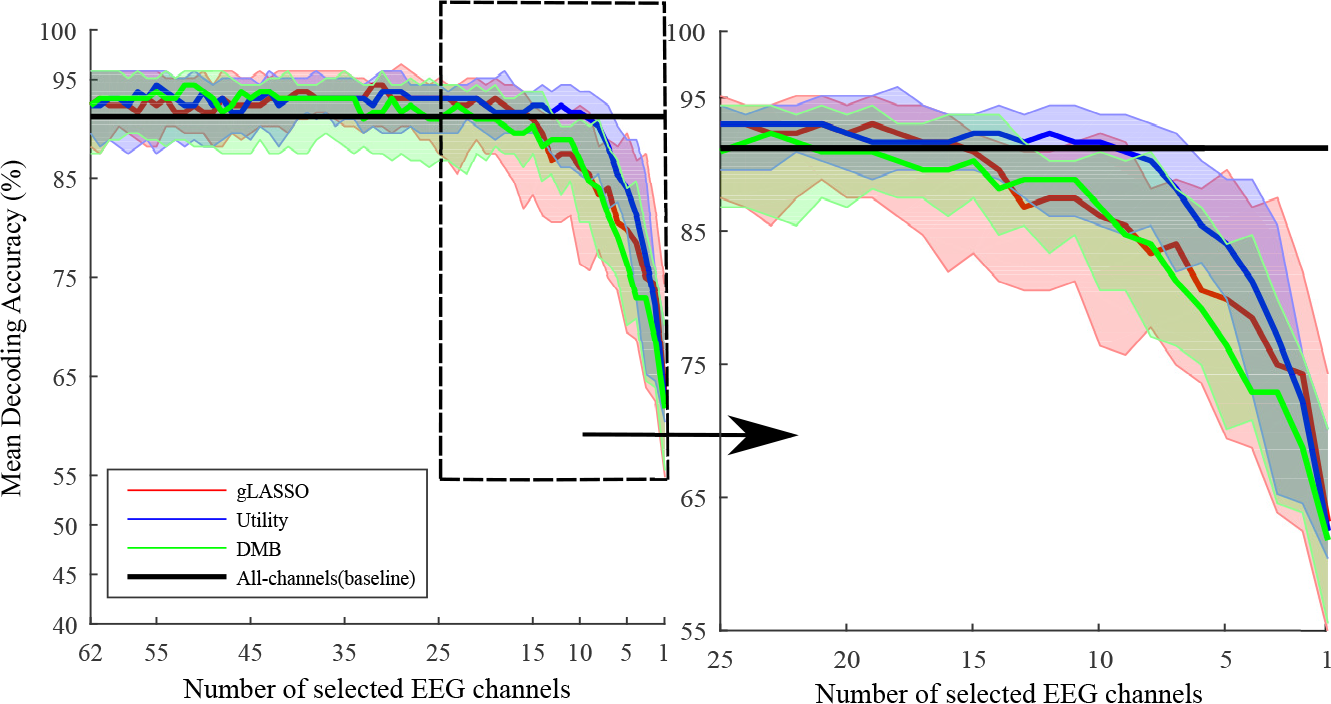
Comparison of three channel selection methods on the Orig (Cz-ref) dataset for subject-dependent channel selection. Median AAD decoding accuracies are plotted as bold lines; The boundaries of the shaded regions form the 25th and 75th percentile accuracies. (a) For number of channels *N* ∈ {1, …, 62} (b) A zoomed plot focusing on *N* < 25.

The *p*-values of the statistical tests are given in Table I (*p*-values are not corrected for multiple comparisons). It can be observed from Table I, in the case of the utility-based method compared to gLASSO, *p*-values are < 0.05 for 4 ≤ *N* ≤ 15 for both the tests. While comparing the utility-based method with DMB, in six out of the eleven values of *N*, *p*– values are < 0.05. For *N* < 4 no significant differences were found. From both Fig. 1 and Table I, a clear trend is observed that the utility-based method outperforms the other methods for small vales of *N*. Therefore, only the utility-based method is used for the optimal node selection problem in the sequel in order to reduce the exploration space.

**TABLE I:**
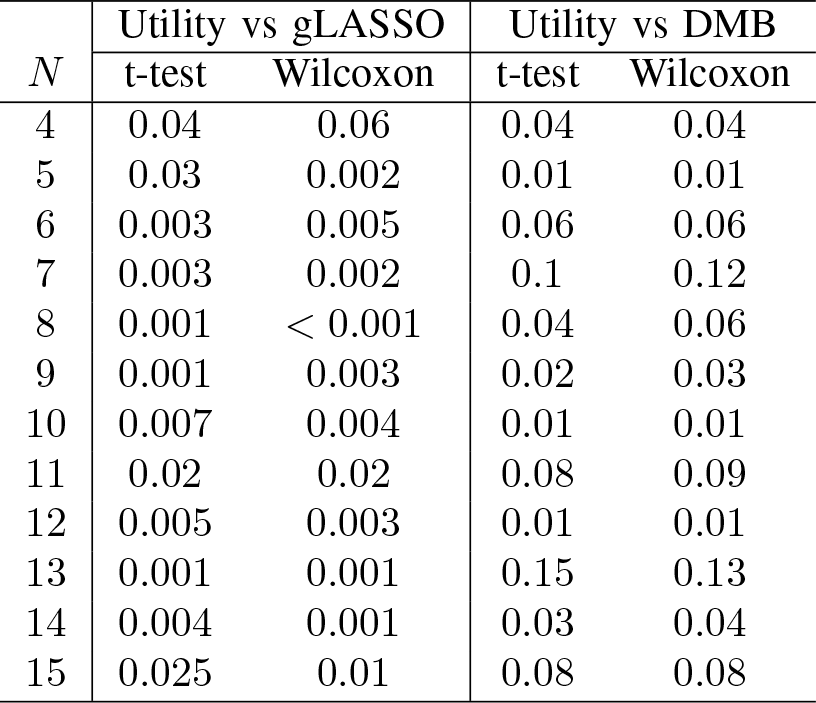
Significance tests comparing channel selection methods: *p*–values obtained using statistical tests comparing utility-based method and both DMB and gLASSO methods for 4 ≤ *N* ≤ 15.

| $N$ | Utility vs gLASSO | | Utility vs DMB | |
| --- | --- | --- | --- | --- |
|  | t-test | Wilcoxon | t-test | Wilcoxon |
| 4 | 0.04 | 0.06 | 0.04 | 0.04 |
| 5 | 0.03 | 0.002 | 0.01 | 0.01 |
| 6 | 0.003 | 0.005 | 0.06 | 0.06 |
| 7 | 0.003 | 0.002 | 0.1 | 0.12 |
| 8 | 0.001 | < 0.001 | 0.04 | 0.06 |
| 9 | 0.001 | 0.003 | 0.02 | 0.03 |
| 10 | 0.007 | 0.004 | 0.01 | 0.01 |
| 11 | 0.02 | 0.02 | 0.08 | 0.09 |
| 12 | 0.005 | 0.003 | 0.01 | 0.01 |
| 13 | 0.001 | 0.001 | 0.15 | 0.13 |
| 14 | 0.004 | 0.001 | 0.03 | 0.04 |
| 15 | 0.025 | 0.01 | 0.08 | 0.08 |

### B. Short-distance node selection

#### 1) Single-channel node selection

In the next experiment, the effect of short-distance EEG measurements is investigated by first applying the utility-based channel selection strategies described in Section II-F to select *N* = 2, 3, … 8 single-channel short-distance nodes from the set *S*_1*ch*_ of all candidate node positions/orientations. Note that the selection of a node in this case corresponds to the selection of a single electrode pair. The AAD performance using these selected nodes is compared with the AAD performance when using the same number of channels from two different long-distance EEG measurement configurations described in Section II-D, viz. Orig (Cz-ref) and Any-Pair.

Fig. 2 shows the result for a subject-dependent node selection. The distribution of the corresponding short-distance node locations is shown in Fig. 3, with Fig. 3a plotting the locations and configurations of the best *N* = 2, 4, 6, 8 nodes and Fig. 3b showing the distribution of electrodes among the selected nodes across all subjects. A paired t-test and a Wilcoxon signed-rank test between the performances using the best short-distance nodes and the best long-distance configurations were carried out and the *p*-values were found to be *>* 0.5 for all *N* in the Cz-ref case, and *>* 0.1 for all *N* (except *N* = 4) in the ‘Any-Pair’ case. For *N* = 4, *p* = 0.04 in the Any-Pair case.

**Fig. 2:**
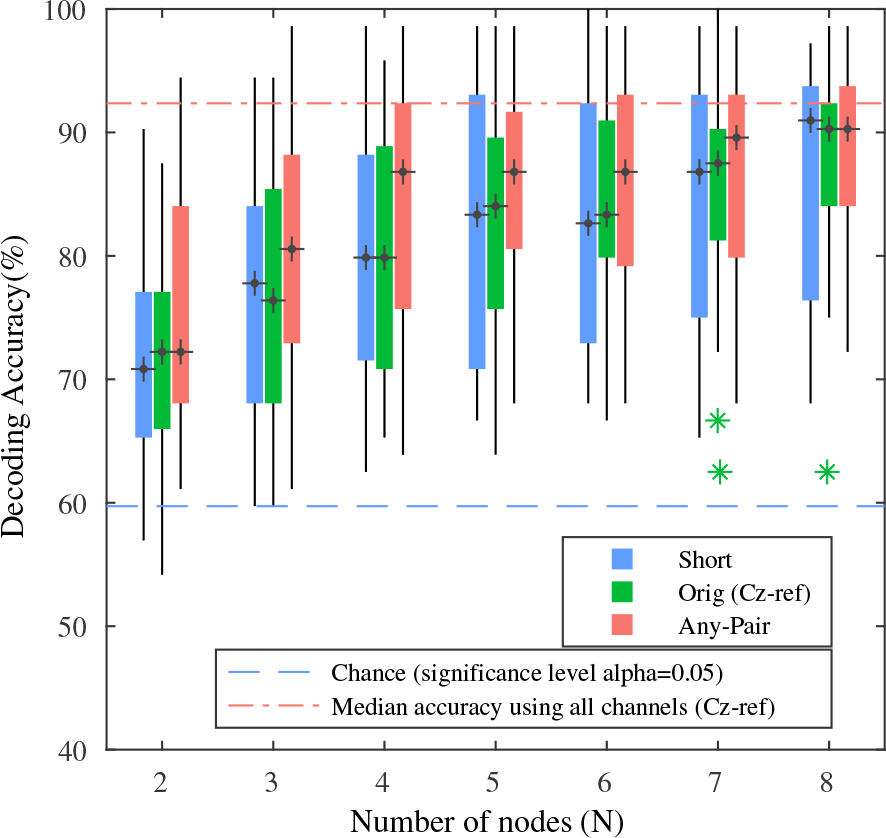
Utility-based subject-dependent node selection: Performance of the *N* best short-distance nodes from *S*_1*ch*_ compared with the *N* best Orig (Cz-ref) and Any-Pair channels. The stars represent outliers which are greater than 1.5 times the interquartile range away from the top or bottom of a box.

**Fig. 3:**
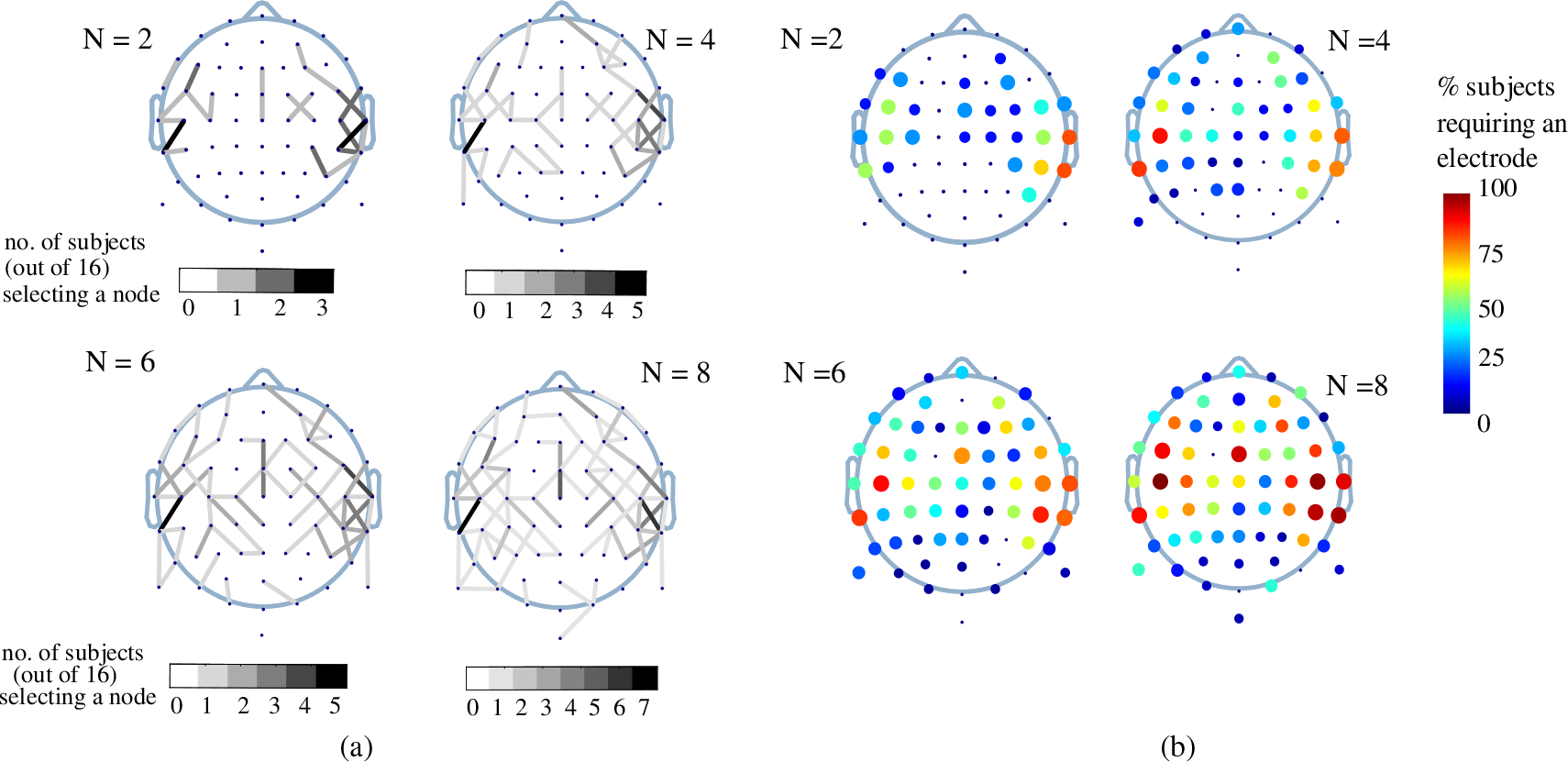
(a) The locations of the best set of short-distance nodes for *N* = 2, 4, 6, 8. The gray-scale of a node indicates the number of subjects which selected that node. (b) Electrodes which are required for the best *N* nodes: Colors of the electrodes indicate the percentage of subjects having a particular electrode in its best *N* nodes. The size of the point representing an electrode is also proportional to the number of times an electrode is present in the best *N* selected nodes.

The best universal single-channel node locations obtained using the utility-based channel selection are shown in Fig. 4. The AAD performance using the universal node selection is compared between short-distance and long-distance recordings in Fig. 5a for a subject-dependent decoder training and in Fig. 5b for a universal decoder training. However, note that in both cases the node *selection* was done in a subject-*independent* fashion.

**Fig. 4:**
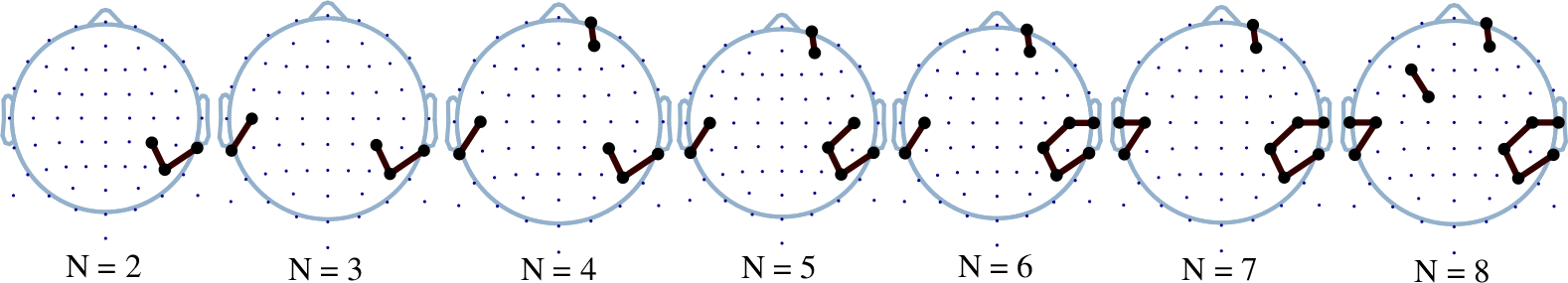
Utility-based universal node selection: Locations of the best universal set of short-distance nodes for *N* = 2, 3, … 8.

**Fig. 5:**
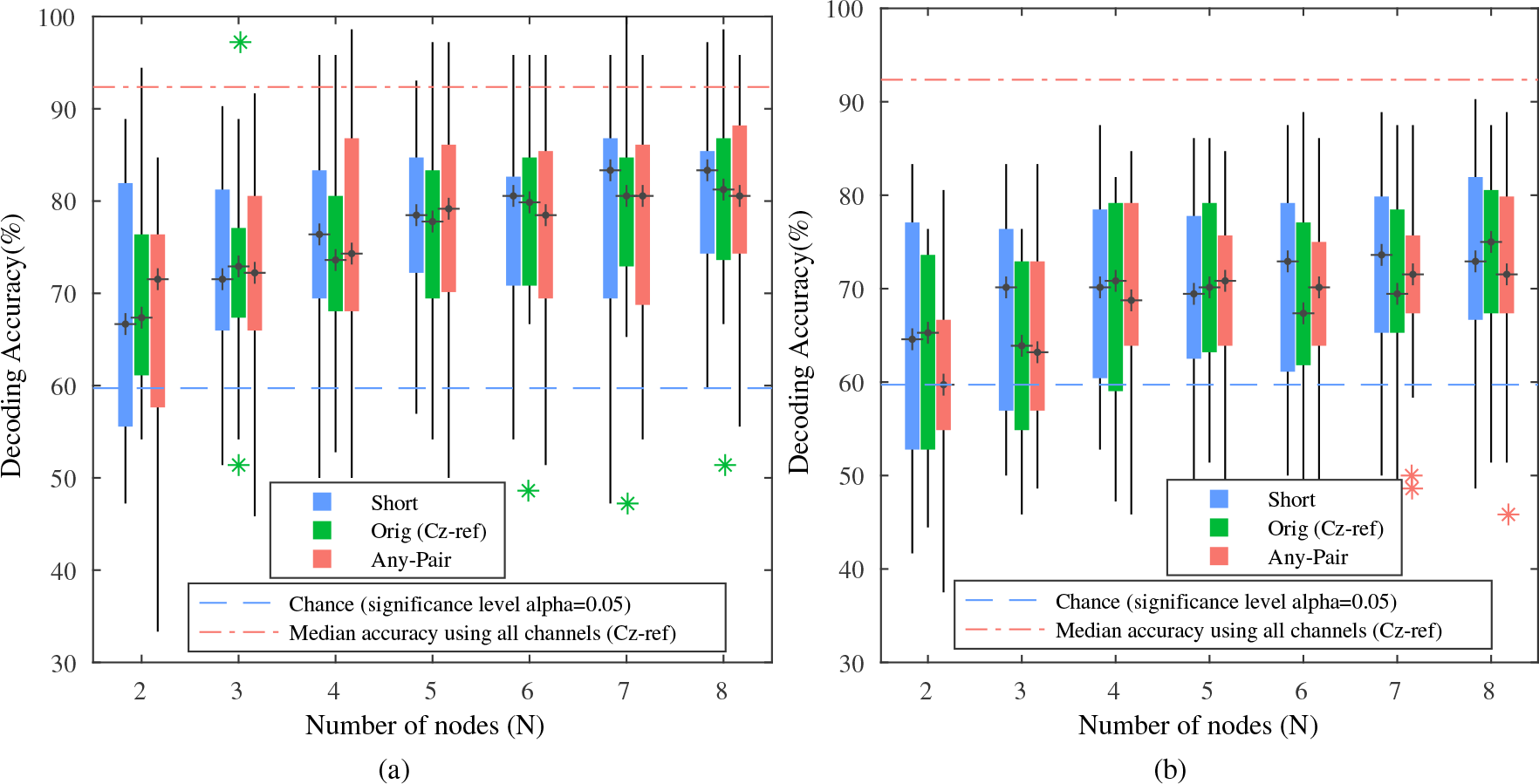
Utility-based universal selection of the *N* = 1,…, 8 best short-distance nodes from *S*_1*ch*_ compared with the *N* best Orig (Cz-ref) and Any-Pair channels: (a) AAD performance for a subject-dependent decoding. (b) AAD performance for a universal decoding. The stars represent outliers which are greater than 1.5 times the interquartile range away from the top or bottom of a box.

Both the t-test and Wilcoxon signed-rank test between universal short-distance node selection and both Cz-ref and Any-pair channel selections, followed by subject-dependent decoding, yielded *p*-values > 0.3 for all values of *N*. When followed by universal decoding, *p*-values were > 0.1 while comparing short-distance nodes and both Cz-ref and Any-pair channels for *N* = 4, 5, 6, 7, 8. Meanwhile, *p*-values were < 0.05 for *N* = 3 when comparing with Cz-ref channels and for *N* = 2, 3 when comparing with Any-ref channels. For these few cases where *p* < 0.05, short-distance nodes are in fact observed to outperform long-distance measurement configurations (also visible in Fig. 5b).

A linear mixed effect model was used to model the relationship between the decoding accuracy using short-distance nodes, the number of nodes and the three scenarios, viz. subject-dependent node selection and decoding, universal node selection followed by subject-dependent decoding and universal node selection followed by universal decoding. In the model, subjects were considered as a random factor. The results showed that the decoding accuracies increase significantly with the number of nodes in all the three scenarios (*p* < 0.001).

#### 2) Two-channel node selection

Next, we investigate the effect of using 2-channel sensor nodes from the set *S*_2*ch*_ instead of single-channel sensor nodes from the set *S*_1*ch*_. The corresponding AAD accuracy is illustrated in Fig. 6 (a) for the subject-dependent node selection. A signed-rank Wilcoxon test and a paired t-test between two-channel and single-channel decoding accuracies resulted in *p*-values < 0.05 for all values of *N* = 1, 2, … 8 for subject-dependent node selection. The results for a universal node selection are shown in Fig. 6(b) and Fig. 6(c) for a subject-dependent decoding and a universal decoding, respectively. Both a signed-rank Wilcoxon test and a paired t-test between two-channel and single-channel decoding accuracies resulted in *p*-values < 0.05 for all values of *N* while using a subject-dependent decoder. However while using a universal decoder, the *p*-values were < 0.05 for only two values of *N*.

**Fig. 6:**
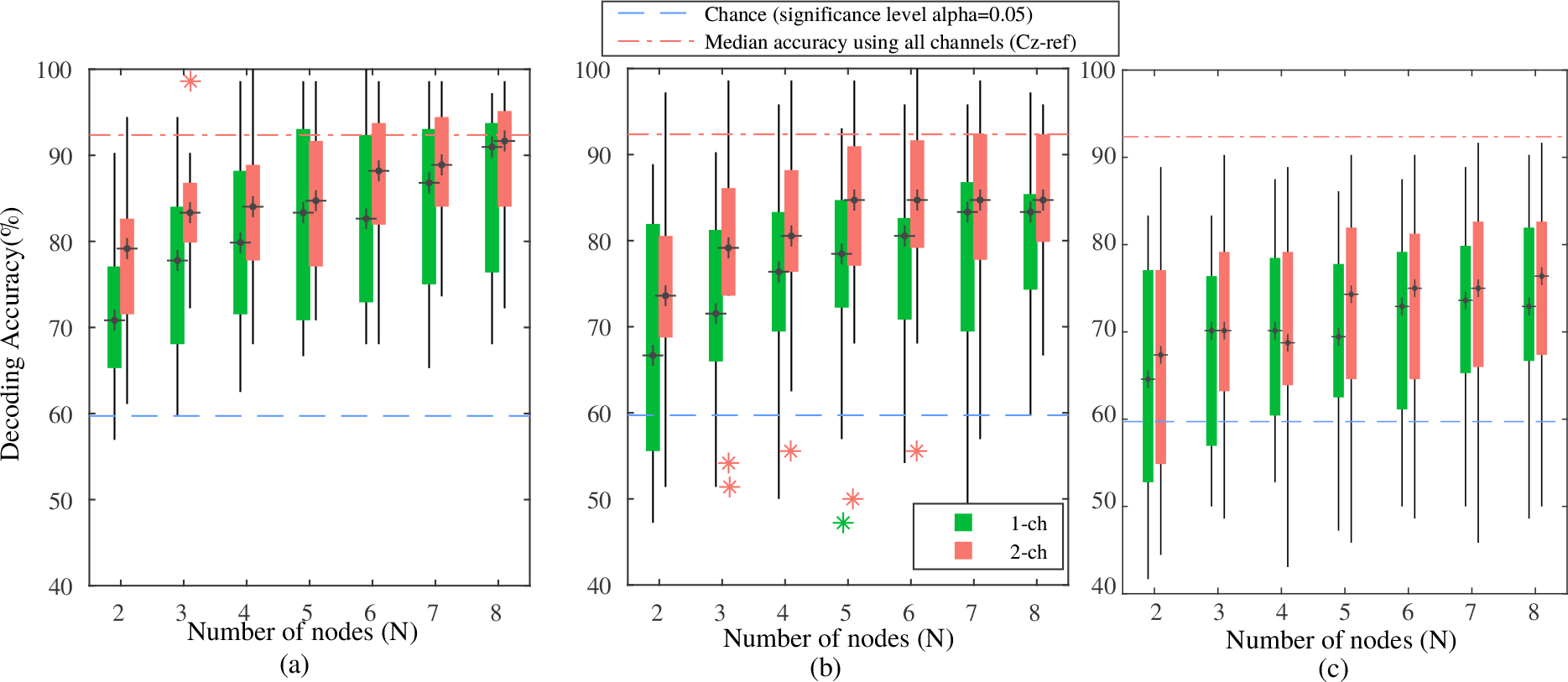
Single-channel vs two-channel nodes: Comparison of decoding performance across subjects for (a) Subject-dependent node selection and subject-dependent decoding (b) Universal node selection and subject-dependent decoding (c) Universal node selection and universal decoding. The stars represent outliers which are greater than 1.5 times the interquartile range away from the top or bottom of a box.

The locations and orientations of the best *N* subject-dependent two-channel nodes are shown in Fig. 7. Similar to the earlier case of single-channel nodes, Fig. 8 shows distribution of the electrodes present in the best *N* two-channel nodes. The locations and orientations of universal two-channel nodes are illustrated in Fig. 9. Note that some of the nodes appear to have electrode pairs that apparently make angles closer to 180 degrees than to 90 degrees. However, this is due to the fact that the electrode positions are mapped from 3D coordinates (on a head) to a 2D plane to visualize them in a topoplot, thereby not preserving the angles.

**Fig. 7:**
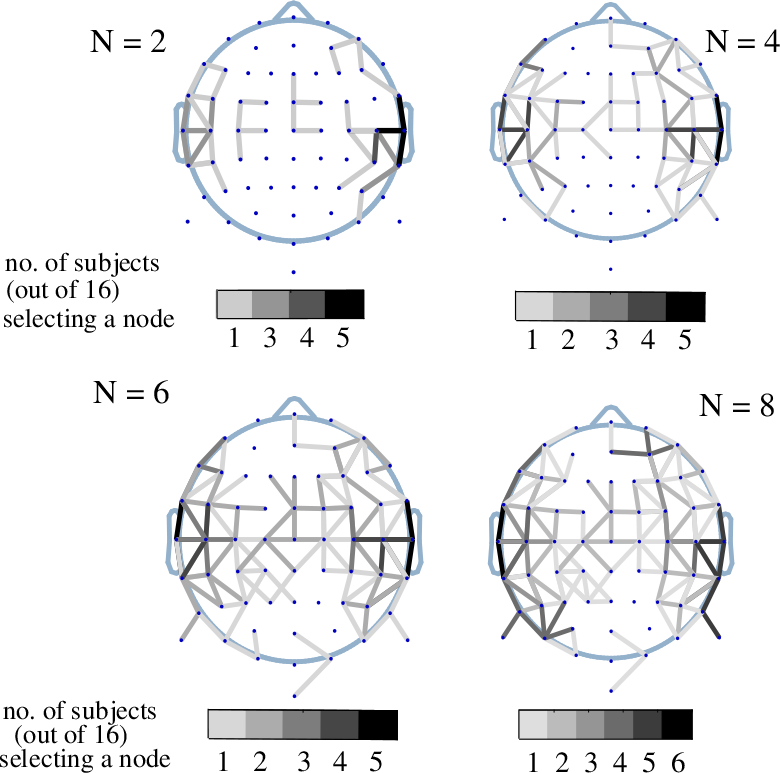
Utility-based two-channel nodes selection: Locations and configurations of the best subject-dependent two-channel nodes.

**Fig. 8:**
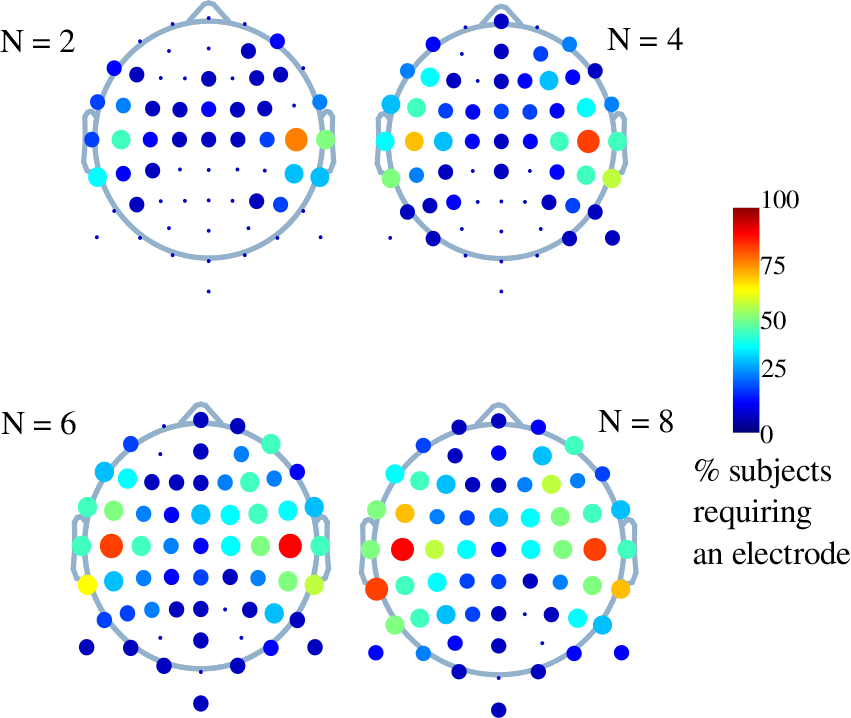
Utility-based two-channel nodes selection: Electrodes which are required for the best *N* nodes.

**Fig. 9:**
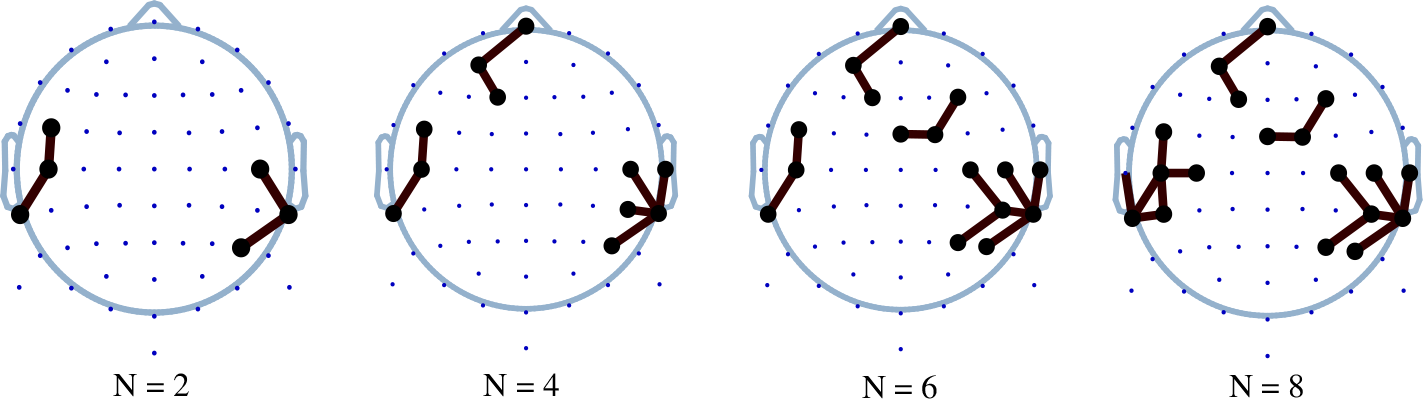
Utility-based selection of two-channel nodes: Locations and configurations of the best universal two-channel nodes.

## IV. Discussion

In Fig. 1, the advantage of the proposed greedy utility-based channel selection over a DMB greedy channel selection strategy [20], [29] and a gLASSO based method [1] was verified, as explained in Section III-A. The figure illustrates the AAD performance using DMB greedy channel selection to be consistent with the results reported in [20] (on a different dataset), where it was concluded that the performance begins to drop significantly when number of channels are reduced to 15 or lower. However, when using a utility-based channel selection strategy, this drop in performance is observed only when the number of channels is lower than 10. Moreover, the figure shows that in general, the utility-based selection selects fewer channels compared to the other two methods to achieve a pre-defined accuracy. The AAD performances using the EEG measurements from the best short-distance nodes were compared to two different long-distance benchmark sets, viz. the Orig (Cz-ref) and Any-Pair channels as described in Section II-D. The results in Fig. 2 suggest that the performances using the best short-distance single-channel nodes were similar to long-distance configurations. Firstly, no significant differences between performance using the best short-distance nodes and the best Cz-referenced channels were found which can be inferred from the *p*-values of the statistical tests detailed in Section III-B1. Secondly, comparing the optimal short-distance measurements with the best Any-Pair channels, no significant differences can be found for all but one value of *N*. In principle, these results only imply that there is not sufficient evidence to reject the null-hypothesis that the performance is the same between all cases, which does not necessarily mean that the null-hypothesis is true. However, all but one *p*-value are far from the *α* = 0.05 significance level, despite the relatively large number of subjects and number of comparisons. As such, there are at least strong indications that the impact of miniaturization is negligible if an equal number of channels are used and if optimal node locations are selected in both cases. This observation is encouraging for the use of concealable and wireless mini-EEG devices and WESNs, where short inter-electrode distances are unavoidable to allow for a sufficient miniaturization of the devices.

In Fig. 3a, the nodes selected by the subjects for *N* = 2, 3, … 8 are predominantly located near the left and the right temporal lobe, where also the auditory cortex is located. This can also be observed in Fig. 3b, where the electrodes located near the same regions are shown to be used most in the node selection. A similar pattern has been reported in the literature for long-distance recordings with a common reference [20]. The optimal universal single-channel node locations plotted in Fig. 4 show a similar trend as the corresponding subject-dependent locations, where also the majority of the nodes are located in the area of the temporal lobe. Note that a few of the nodes in this figure and Fig. 3a share electrodes. In practice mini-EEG devices could instead be placed close to each other with positions close to the selected ‘common’ electrode position on the scalp. If necessary, it can be avoided to obtain a selection where some of the nodes share electrodes by applying a restart procedure for such collisions as in [1]. However, it was demonstrated in [1] that this does not significantly affect AAD performance, while it can lead to excessively long computation times.

Two different approaches were used to obtain the AAD decoding accuracies with universal locations, which are shown in Fig. 5a and Fig. 5b. The results again demonstrate that short-distance nodes do not yield a significantly lower AAD performance compared to using the same number of long-distance EEG measurements. Indeed, no statistically significant improvement were found for long-distance configurations over short-distance nodes with both a paired t-test and Wilcoxon signed-rank test as detailed in Section III-B1. The Fig. 5b, which plots decoding accuracies obtained using a universal decoder, also indicate that short-distance measurements deliver at least the same performance as long-distance measurements. A linear mixed effects modeling of the performance with respect to the number of nodes, which is detailed in Section III-B1, showed that there is a significant effect of the number of short-distance nodes *N* on the decoding accuracies, i.e., increasing the number of mini-EEG devices leads to an improvement in AAD performance. For both subject-dependent and universal node selection.

We also investigated the effect of adding a third electrode in each node, yielding two EEG channels. Since both of these channels are recorded by electrodes that are very close to each other (within the same node), it is a-priori not obvious whether the extra channel is useful or whether it is mostly redundant. Fig. 6 suggests that, adding this additional channel does have a clear benefit. In Fig. 6 (a), two-channel nodes can be observed to perform better than the same number of single-channel nodes in the case of subject-dependent node selection and decoding. This observation can also be noted in Fig. 6 (b) for universal node selection followed by subject-dependent decoding. The *p*-values of the statistical tests comparing the two performances reported in Section III-B2 support this observation in the two cases. These results show that even at short inter-electrode distances, the extra channel can cause significant improvement in performance despite the fact that all three electrode are close to each other. This may be due to the almost-orthogonal orientation of the two electrode pairs, which provides two orthogonal axes (instead of only one) to capture dipoles in all directions parallel to the plane spanned by the electrode pair, as opposed to a single direction in the case of 1-channel nodes. It should be noted that a dipole orthogonal to the plane spanned by the electrode pair will still not be captured, although it may be captured by another node on a different scalp position that better aligns with this orientation. In the case of a universal decoder (see Fig. 6 (c)), the beneficial effect of adding this extra channel to each node is less clear, which implies that the extra orientation can only be properly exploited by a subject-dependent decoder. Statistical tests performed between the two cases do not confirm a significant benefit for two-channel nodes in this case. This could be explained by the fact that the relevant dipole orientations may differ from subject to subject, such that the extra degree of freedom added by the extra channel to tune the decoder to particular dipole orientations cannot be exploited by a universal decoder, as it cannot adapt its weights to the individual subject.

The orientations of the best two-channel nodes and the distribution of electrodes in these nodes are shown in Fig. 7 and Fig. 8 respectively for subject-dependent node selection. Fig. 9 shows the location and orientations of universal node selection. It should be noted here that the best two-channel nodes are selected from a candidate set of two-channel nodes as explained in Section II-C and that they are not derived from best single-channel nodes. Nevertheless, it can be observed from Fig. 7, Fig. 8 and Fig. 9 that the best two-channel nodes are again mostly located close to the auditory cortex within the temporal lobe.

## V. Conclusion

Miniaturized EEG (mini-EEG) sensor devices are becoming increasingly prevalent in the field of neural signal processing and pave the way towards chronic neuromonitoring applications. Hence, it is essential to understand the effects of this miniaturization on EEG signal processing methods which have been tested with traditional EEG equipment. In this work, the effect of short-distance EEG measurements that arise with miniaturization of EEG sensor devices was investigated within the context of an AAD task, which may be used in future-generation neuro-steered auditory prostheses, e.g., for the cognitive control of hearing aids or cochlear implants [23], [24].

We have shown that short-distance referencing in a mini-EEG device has little impact on AAD performance when compared to the commonly used Cz-referencing and an any-reference long-distance referencing configuration, provided the electrodes of the mini-EEG devices are placed at ideal locations. These results are encouraging for the use of multiple mini-EEG devices as nodes in a WESN to perform chronic neuromonitoring for AAD. We have also proposed a utility-based greedy channel selection strategy for the (redundant) channel selection problem in AAD, which outperforms two other channel selection methods, and which was used to select the ideal locations for placing mini-EEG devices. We have found that these locations are close to the auditory cortex within the temporal lobe, which is consistent with previous results found in the literature. We have also shown that having two-electrode pairs in a mini-EEG node results in a significant improvement in AAD performance over single-channel nodes even at short inter-electrode distances, but only when a subject-dependent decoder can be trained, in order to exploit the additional degree of freedom to capture relevant dipoles with any orientation within each subject.

## VI. Acknowledgement

The authors would like to thank the lab ExpORL (KU Leuven), and in particular Prof. T. Francart, N. Das and W. Biesmans for sharing their auditory attention EEG data set.

1 For example, scaling a channel with a factor *x* will scale the corresponding LS decoder weight with a factor 1/*x*, while the information provided by the signal remains the same.

2 Since the original 64 channel EEG data was rereferenced to the Cz electrode, the total effective number of channels reduce to 63.

